# Measuring the Impact of an Open Online Prescribing Data Analysis Service on Clinical Practice: a Cohort Study in NHS England Data

**DOI:** 10.1101/306415

**Authors:** Alex J Walker, Helen J Curtis, Richard Croker, Seb Bacon, Ben Goldacre

**Affiliations:** Evidence Based Medicine DataLab, Nuffield Department of Primary Care Health Sciences, University of Oxford, Radcliffe Observatory Quarter, Woodstock Road, Oxford OX2 6GG

**Keywords:** Drug Prescribing, Cost Control, Patient Safety, Treatment Efficacy

## Abstract

**Background:** OpenPrescribing is a freely accessible service that enables any user to view and analyse NHS primary care prescribing data at the level of individual practices. This tool is intended to improve the quality, safety, and cost-effectiveness of prescribing.

**Objectives:** We set out to measure the impact of OpenPrescribing being viewed on subsequent prescribing.

**Methods:** Having pre-registered our protocol and code, we measured three different metrics of prescribing quality (mean percentile across 34 existing OpenPrescribing quality measures, available “price-per-unit” savings, and total “low-priority prescribing” spend) to see if they changed after CCG and practice pages were viewed. We also measured whether practices whose data were viewed on OpenPrescribing differed in prescribing, prior to viewing, to those who were not. We used fixed effects and between effects linear panel regression, to isolate change over time and differences between practices respectively. We adjusted for month of prescribing in the fixed effects model, to remove underlying trends in outcome measures.

**Results:** We found a reduction in available price-per-unit savings for both practices and CCGs after their pages were viewed. The saving was greater at the practice level (−£40.42 per thousand patients per month, 95% confidence interval −54.04 to −26.01) than at CCG level (−£14.70 per thousand patients per month, 95% confidence interval −25.56 to −3.84). We estimate a total saving since launch of £243k at practice level and £1.47m at CCG level between the feature launch and end of follow-up (August to November 2017) among practices viewed. If the observed savings from practices viewed were extrapolated to all practices, this would generate £26.8m in annual savings for the NHS, approximately 20% of the total possible savings from this method. The other two measures were not different after CCGs/practices were viewed. Practices which were viewed had worse prescribing quality scores overall, prior to viewing.

**Conclusions:** We found a clinically significant positive impact from use of OpenPrescribing, specifically for the class of savings opportunities that can only be identified by using this tool. We also show that it is possible to conduct a robust analysis of the impact of such an online service on clinical practice.

## Background

The OpenPrescribing project aims to make prescribing data more accessible and impactful for clinicians, policy makers and others. It does this through providing a user friendly web interface. It is hoped that this will enable safer, more effective and cost efficient prescribing by making users more aware of their prescribing behaviour, by making comparisons with peers and by highlighting meaningful changes over time. OpenPrescribing gives a range of specific analyses and tools for each CCG and practice in three broad classes set out in Box 1.

### Box 1. Prescribing Data Available on OpenPrescribing

Prescribing data analyses on OpenPrescribing for individual institutions include:

1. A range of prespecified prescribing measures (for example “proportion of statins that are high-cost”, “antibiotics prescriptions per standardised population unit”).
2. A “Price Per Unit” tool that uses a novel method to identify a range of newly identified cost-saving opportunities bespoke for each institution, driven by analysis of national price variation within the same chemical and dose across each month. Savings are identified by very-large-scale data analysis across the entire national dataset and at present, crucially for the analysis presented here, these savings can only be identified using OpenPrescribing[6]. Savings are best realised by viewing practice-level data.
3. A range of tools that collectively calculate the total spend on items identified by NHS England as “low priority” on grounds of extremely low cost-effectiveness [7].

There are many commercial tools that aim to improve prescribing in the UK. These include Oracle [1], Optum (ScriptSwitch) [2], Prescribing Services [3], and Prescribing Support Services [4]. While these tools may be effective at generating change, there is little publicly available evidence of robust testing. Such testing is a important element of good commissioning practice, in order to ensure that resources are used cost-effectively [5].

We set out to deliver a robust quantitative evaluation of the impact of use of an open online tool, that may act as a template for other evaluations of similar tools, both in terms of the open and reproducible approach, and the methodology used. We set out to measure whether any change in prescribing behaviour occurs after use of OpenPrescribing, using the most robust observational methods to determine whether there is any causal association. Because it is possible that practices and CCGs who engage with their prescribing data are systematically different to those who don’t, we also set out to measure whether practices and CCGs viewing OpenPrescribing already differ from their peers in prescribing behaviour, prior to using the tool.

## Methods

### Prespecification and protocol registration

As this is an observational study of the impact of our own service we endeavoured to minimise any potential for conflict of interest impacting on results by fully prespecifying our methods and posting the protocol on the Open Science Framework prior to commencement [8] (osf.io/9aj53). In the protocol we specify the outcomes to be measured, along with the full analytic approach. The entire analytic code was written against a small sample of seven institutions’ data, and also published prior to conducting the analysis. There were no substantial changes to the outcomes or methodology (including analytic code) between prespecification and reporting the results here.

### Data and Sources

Prescribing outcome data were obtained from the monthly prescribing dataset published by NHS digital and aggregated by the OpenPrescibing project. The monthly prescribing datasets contain one row for each treatment and dose, in each prescribing organisation in NHS primary care in England, describing the number of prescriptions issued and the total cost. Each practice in England belongs to one of 207 Clinical Commissioning Groups (CCGs); we aggregated practice data to these CCGs for CCG-level analyses. Practices with a very small list size <1000 were excluded due to likelihood of being an atypical practice.

Data on the exposure (CCG and practice page views) were obtained from Google Analytics for OpenPrescribing.net. We collected page view data from the launch of the project (1st December 2015) to the most recent data available at the time of extraction (14th Jan 2018). Page view data contains the date of each view, along with which practice or CCG’s data were viewed, and whether any specific site features were viewed (e.g. the price-per-unit or low-priority features). Page view dates were aggregated to month-year.

### Exposures

The exposures used for this study are pageviews on the OpenPrescribing.net site. This is a proxy exposure, as we are not able to attribute site use to an individual practice/CCG, but instead assume that a high proportion of traffic to each practice/CCG’s prescribing page is from persons associated with that practice or CCG.

We generated two different exposure variables to determine the associations with viewing OpenPrescribing. Firstly, for each month, in each practice and CCG, we defined whether this was ‘before viewed’, during ‘month first viewed’, or ‘after first view’ the first view of data on OpenPrescribing. The ‘first view’ is defined as the first month that a practice or CCG page was viewed more than once (in order to exclude one-off visits, which are unlikely to represent full engagement with the site). Secondly, we generated a variable to describe the total number of views of each practice and CCG page on OpenPrescribing (divided into categories: 0 views; then tertiles of number of practice views amongst those who were viewed).

For the price-per-unit and low-priority outcomes (described below) the exposure was restricted to views of the price-per-unit and low-priority pages for an institution (for example https://openprescribing.net/ccg/99P/price_per_unit/).

### Outcomes

We used three outcome variables in order to measure the effectiveness of the three main features of OpenPrescribing (see Box 1). Firstly, we calculated mean percentile for each of the standard prespecified OpenPrescribing measures [9], excluding the NHS England low-priority measures (which are analysed separately below) and those where a value judgement is not made (currently direct acting oral anticoagulants [10] and two pregabalin measures [11,12]). OpenPrescribing measure performance was aggregated by taking the mean percentile across all included measures for each practice or CCG in each month. Secondly, price-per-unit efficiency was calculated as the total identified price-per-unit savings available for each practice and CCG in each month: full methods for calculating “price-per-unit” savings are described elsewhere [6]. Thirdly, we calculated total spending on NHS England “low-priority” measures (as described elsewhere [7]) for each practice and CCG in each month.

Outcomes were measured on a monthly basis, over a time period from three months before the launch of the respective tool (to obtain a suitable baseline) to the most recent available data. Launch dates for the outcomes were December 2015 for the OpenPrescribing measures (i.e. the OpenPrescribing service as a whole), August 2017 for the price-per-unit feature, and September 2017 for the low-priority spend feature.

### Analysis

Our analysis is described in detail in our prespecified analytic code which is shared in full [8]. Analyses were performed separately at practice level and CCG level. We used fixed effects panel regression and between effects panel regression in order to limit measurement of variation within practice or CCG (i.e. variation over time), and between practice or CCG, respectively.

#### CCG and practice views

In addition to the analysis prespecified in the protocol [8], we calculated summary statistics for the number of practices and CCGs being viewed for each outcome, in order to provide further context to the analysis.

#### Before and After Viewing

To measure change in prescribing outcomes over time (within CCG/practice effects), we used fixed effects linear panel regression, to remove the effect of time-invariant (between practice) characteristics. We firstly used a univariable model, then added calendar month to the model, in order to adjust for underlying national changes over time. This should leave only differences over time between practices that have and have not been viewed on OpenPrescribing.

#### Before Viewing

To measure differences between practices that have and have not been viewed on OpenPrescribing, we used between effects linear panel regression. This was a simple univariable model to test the hypothesis, with the between effects model removing any effects occurring over time. In order to remove any potential influence of OpenPrescribing, we used the the three month period prior to the above launch dates for each outcome (as described in the outcomes section).

## Results

### CCG and practice views

Of 207 CCGs included in the study, 207 (100%) were counted as exposed (≥2 views in the same month) for the mean measure outcome during at least one month, while 127 (61.4%) were viewed for the price-per-unit outcome, and 68 (32.9%) for the low-priority prescribing outcome. 7,318 practices were included in the study: of these 4,578 (62.6%) were viewed in at least one month for the mean measure outcome, 279 (3.8%) for the price-per-unit outcome, and 59 (0.8%) for the low-priority outcome.

### Prescribing before and after viewing OpenPrescribing

Table 1 shows the change in prescribing outcomes measured at CCG level, during and after each CCG was first viewed on OpenPrescribing. Table 2 shows the same but at practice level. Univariable results include secular trends that exist regardless of any influence of OpenPrescribing: this crude unadjusted data is presented only for reference; multivariable results account for secular trends and show the impact of OpenPrescribing views. There was no change in mean OpenPrescribing measure score at either CCG or practice level. Although there is a significant change at practice level in the univariable analysis, this effect was eliminated by adjusting for calendar month.

There was a reduction in available price-per-unit savings after CCGs (Table 1) and practice (Table 2) were viewed on OpenPrescribing. The effect size was greater at practice level (−£40.42 per thousand patients per month, 95% confidence interval, −54.04 to −26.01) than at CCG level (−£14.70 per thousand patients per month, 95% confidence interval, −25.56 to −3.84). In the univariable analysis there was a much greater effect size due to the overall trend of decreasing available savings over the study period, but the effect remained after adjustment for calendar month.

Multiplying the estimated (per thousand patient) saving by the CCG and practice populations in the “after looking” months gives a total estimated saving of £1.47m (95% confidence interval, £384k to £2.56m) at the CCG level and £243k (95% confidence interval, £162k to £326k) at the practice level, in practices and CCGs whose data was viewed. It is possible that some of these savings will overlap and therefore it is not appropriate to add the two figures together to create total savings. Extrapolating these savings figures to all CCGs and practices across England, if all institutions’ data was viewed, would generate an estimated annual saving of £9.7m at CCG level and £26.8m at practice level.

Total ‘available’ savings calculated by the tool for the time after CCGs/practices were viewed were £31.3m at CCG level £2.4m at practice level. In our paper [6], we estimated that around half of these ‘available’ savings might be ‘achievable’. This means that around 10% of the ‘achievable’ savings were realised at CCG level, and around 20% realised at practice level.

There was no change in the total spend on low-priority measures at CCG or practice level. The small reduction seen at CCG level in the univariable analysis is again eliminated after adjustment for calendar month.

**Table 1:** CCG level results of fixed effects linear panel regression, showing the change in each outcome before and after the corresponding CCG page on OpenPrescribing.net was viewed.

| Outcome | Time period | Mean value<br>(higher is worse) | Univariable |  |  | Adjusted for calendar month |  |  |
| --- | --- | --- | --- | --- | --- | --- | --- | --- |
|  |  |  | Change | 95% CI |  | Change | 95% CI |  |
| Mean measure score | Not viewed | 50.1% | Reference |  |  | Reference |  |  |
|  | Month first viewed | 50.0% | 0.00 | -0.24 | 0.23 | -0.09 | -0.42 | 0.23 |
|  | After first view | 49.9% | -0.05 | -0.14 | 0.04 | -0.26 | -0.60 | 0.08 |
| Price-per-unit | Not viewed | £351.19 | Reference |  |  | Reference |  |  |
|  | Month first viewed | £335.94 | -£78.28 | -£92.32 | -£64.24 | -£10.30 | -£21.87 | £1.26 |
|  | After first view | £283.72 | -£132.75 | -£143.45 | -£122.06 | -£14.70 | -£25.56 | -£3.84 |
| Low-priority measures | Not viewed | £191.68 | Reference |  |  | Reference |  |  |
|  | Month first viewed | £192.79 | -£11.13 | -£15.33 | -£6.93 | -£2.13 | -£6.41 | £2.15 |
|  | After first view | £194.51 | -£9.94 | -£14.59 | -£5.29 | £1.08 | -£3.91 | £6.07 |

**Table 2:** Practice level results of fixed effects linear panel regression, showing the change in each outcome before and after the corresponding practice page on OpenPrescribing.net was viewed.

| Outcome | Time period | Mean value<br>(higher is worse) | Univariable |  |  | Adjusted for calendar month |  |  |
| --- | --- | --- | --- | --- | --- | --- | --- | --- |
|  |  |  | Change | 95% CI |  | Change | 95% CI |  |
| Mean measure score | Not viewed | 45.8% | Reference |  |  | Reference |  |  |
|  | Month first viewed | 46.0% | -0.35 | -0.46 | -0.24 | -0.07 | -0.18 | 0.05 |
|  | After first view | 46.0% | -0.52 | -0.57 | -0.47 | -0.04 | -0.10 | 0.02 |
| Price-per-unit | Not viewed | £426.86 | Reference |  |  | Reference |  |  |
|  | Month first viewed | £419.51 | -£98.57 | -£116.96 | -£80.18 | -£31.49 | -£48.12 | -£14.86 |
|  | After first view | £385.70 | -£148.36 | -£163.26 | -£133.46 | -£40.42 | -£54.04 | -£26.81 |
| Low-priority measures | Not viewed | £191.68 | Reference |  |  | Reference |  |  |
|  | Month first viewed | £192.79 | -£12.84 | -£34.87 | £9.18 | -£4.33 | -£26.32 | £17.66 |
|  | After first view | £194.51 | -£13.15 | -£41.05 | £14.74 | -£3.11 | -£30.97 | £24.76 |

### Pre-existing differences between practices that have/have not been viewed

Table 3 shows the differences in prescribing outcomes before each service was launched, at CCG level, according to the level of views for each CCG. Table 4 shows the same at practice level. CCGs that have been viewed had higher available price-per-unit savings (that is, they were less cost-efficient as prescribers), but similar for the other two outcome measures. For practices, those that have been viewed were worse for price-per-unit and low-priority spending, but similar with respect to mean measure score.

**Table 3:** CCG level results of between effects linear panel regression, showing the differences in in each outcome between CCG pages that were viewed on OpenPrescribing.net at various levels and those that were not.

| Outcome | Number of views | Mean value (higher is worse) | Change | 95% CI |  |
| --- | --- | --- | --- | --- | --- |
| Mean measure score | 0 | No CCGs have 0 views |  |  |  |
|  | 32-135 | 50.6% | Reference |  |  |
|  | 139-228 | 50.4% | -0.20 | -2.93 | 2.52 |
|  | 231-2219 | 48.9% | -1.74 | -4.50 | 1.01 |
| Price-per-unit | 0 | £365.67 | Reference |  |  |
|  | 1-3 | £366.31 | £0.64 | -£61.16 | £62.43 |
|  | 4-9 | £416.62 | £50.95 | -£15.40 | £117.29 |
|  | 10-137 | £444.80 | £79.12 | £14.79 | £143.46 |
| Low-priority measures | 0 | £198.19 | Reference |  |  |
|  | 1 | £200.15 | £1.96 | -£20.40 | £24.32 |
|  | 2-4 | £193.84 | -£4.36 | -£27.85 | £19.14 |
|  | 5-52 | £196.76 | -£1.44 | -£24.80 | £21.93 |

**Table 4:** Practice level results of between effects linear panel regression, showing the differences in each outcome between practice pages that were viewed on OpenPrescribing.net at various levels and those that were not.

| Outcome | Number of views | Mean value (higher is worse) | Change | 95% CI |  |
| --- | --- | --- | --- | --- | --- |
| Mean measure score | 0 | 45.6% | Reference |  |  |
|  | 1-4 | 45.7% | 0.11 | -0.76 | 0.99 |
|  | 5-8 | 46.1% | 0.49 | -0.41 | 1.39 |
|  | 9-343 | 46.7% | 1.21 | 0.32 | 2.10 |
| Price-per-unit | 0 | £477.29 | Reference |  |  |
|  | 1 | £480.02 | £3.95 | -£24.62 | £32.52 |
|  | 2 | £544.99 | £69.42 | £23.84 | £114.99 |
|  | 3-49 | £524.52 | £47.59 | £13.38 | £81.81 |
| Low-priority measures | 0 | £187.72 | Reference |  |  |
|  | 1 | £229.80 | £41.88 | £25.42 | £58.35 |
|  | 2-26 | £251.37 | £63.94 | £37.81 | £90.08 |
|  | Insufficient views for additional tertile |  |  |  |  |

## Discussion

### Summary

We found that the total available price-per-unit savings decreased following views of OpenPrescribing. This saving corresponds to a total measured decrease in spend of £243k at practice level, and £1.47m at CCG level, between the feature launch and end of follow-up (August to November 2017). Our analytic methods make every effort to remove confounding effects, such as differences between institutions, or national secular trends. Extrapolating the observed savings nationally would generate a total saving of £9.7m per year at CCG level and £26.8m per year at practice level. Savings from this new tool were calculated with only one to three months of follow-up data, and may increase over time. We did not find any changes in the overall prescribing score or low-priority measure spend; possible reasons are discussed below. Contrary to expectations, we found that institutions whose data was viewed on OpenPrescribing were overall performing more poorly on prescribing measures prior to viewing.

### Strengths and Weaknesses

We were able to remove the effect of between practice (time-invariant) confounding effects, with the use of fixed effects linear panel regression. This meant that only differences occurring over time, before and after viewing, were measured. In addition to this, adjusting for calendar month allowed us to appropriately remove the effect of any overall national trends over time, which are independent of any effect that OpenPrescribing might have. By fully prespecifying our methods, and publishing our protocol and analytic code in advance of conducting our analysis, we reduce as far as possible the impact of any potential conflict of interest.

We were only able to use a proxy indicator of each institution’s use of the OpenPrescribing tools, since we cannot determine who exactly is viewing the web site, only that a given institution’s pages have been viewed. This is likely to have added noise to the data and, as a result, is likely to have reduced our ability to detect any impact from the tools, as persons not associated with a practice or CCG can view the site (meaning an institution is incorrectly counted as exposed) but cannot impact on prescribing choices. It is difficult to estimate how great this effect might be.

The number of views for each feature on the site varied substantially, largely because newer tools (such as price per unit, and low priority measures) have only been available for a short period of time. This means it was only possible to measure the effects on a relatively small proportion of all practices for these tools. In addition, we specified the start time for possible impact from the standard prespecified prescribing measures as December 2015 because this is when the OpenPrescribing site launched; however, very few measures were available at initial site launch, and these were added gradually over the following two years. This substantially reduced our ability to detect an impact from the standard prescribing measures; however there is no methodologically straightforward means to account for this variation in the characteristics of the exposure over time.

### Findings in Context

There are many providers of services related to medicines optimisation [1–4]. Such providers make varying degrees of claims, with some just being a description of the services provided, while others make strong claims of efficacy. For example, ScriptSwitch claims to have “delivered over £50m in savings to the NHS” [2], but it is not obvious how this was measured, nor over what time period these savings were made. Another example is from Oracle, which claims that it has “enabled antibiotic prescribing to be reduced by 7% [13,14]: however we are aware of no evidence being given for this very substantial claim, which must be interpreted in the context of a pre-existing downward trend in antibiotic prescribing, and extensive work to reduce antibiotic prescribing following the Chief Medical Officer’s prioritisation of the issue in 2013 [15,16]. A notable exception is Prescribing Services, which makes some attempt to measure the impact of its “Risk Stratification in Prescribing & Screening” tool [17] and claims to have reduced emergency admissions: however the methodology used is not described in any detail.

### Policy implications and interpretation

We found a clinically significant positive impact from use of our prescribing data tool. We also show that it is possible to conduct a robust analysis of the impact of such a tool. We will continue to monitor the effectiveness of existing and new features, in order to refine the tools and monitor impact.

We found the price-per-unit savings to be higher per patient at practice level than CCG level: this is to be expected as tailored prescribing changes from this tool are best identified at the level of individual practices; as discussed in prior work [6]. However, these savings might be achieved more simply and efficiently through national policy change.

The lack of positive effect on the overall prescribing measure might be explained in part by the construction of the mean score. There are 34 different measures making up the mean score, so improvements made by a practice or CCG focusing on one measure in particular is likely to be drowned out by noise of other measures. While we would like to have measured the impact on each measure individually, this would make the analysis extremely complex due to the variety between measures, and would also make interpretation more difficult. It would have been preferable to use a more parametric method to summarise the measures, such as mean Z-score, but this was not possible due to non-normal distributions (e.g. bimodal). Additionally, in contrast with the price-per-unit outcome, where OpenPrescribing is the only known source of determining such savings, it is possible that many practices and CCGs are already aware of many of the issues and had existing work to improve prescribing independently of OpenPrescribing use. This is true for many of the standard OpenPrescribing measures; and for the NHS England low-priority measures, which obtained some publicity when they were announced. Other possible reasons for the lack of effect with the low-priority outcome are the potential for changes in one type of prescribing to be lost in the noise of the others, and the lack of follow-up time since launch (at most three months).

Prior to this analysis we hypothesized that practices/CCGs that looked at OpenPrescribing might already have better overall prescribing than those who do not, on the grounds that institutions who are pro-actively engaged with their data are likely to also be more effective at improving their prescribing. In fact we found that the opposite is true for some outcomes. However, this may reflect the way the OpenPrescribing site operates, in that various features highlight practices that are performing the least well on specific prescribing measures: for example, when examining the performance on one measure for all practices in a CCG, practices are ordered from worst to best, increasing the likelihood that worse practices will be clicked on when browsing. Similarly, the OpenPrescribing “email alerts” service uses various statistical process control methods to highlight specific practices and CCGs with worse prescribing.

## Conclusions

We found a clinically significant positive impact from use of our prescribing data tool. We also show that it is possible to conduct a robust analysis of the impact of such a tool. We will continue to monitor the performance of the OpenPrescribing services as more follow-up time accrues, and as features are added and enhanced. Our methods, and full prespecification, may represent a good template for similar impact assessments on services which aim to improve healthcare.

